# The growth and expansion of meningeal lymphatic networks are affected in craniosynostosis

**DOI:** 10.1101/2021.08.27.457986

**Authors:** Phillip Ang, Matt Matrongolo, Max A Tischfield

## Abstract

Congenital skull malformations are associated with vascular anomalies that can impair fluid balance in the central nervous system. We previously reported that humans with craniosynostosis and mutations in *TWIST1* have dural venous sinus malformations. It is still unknown whether meningeal lymphatic networks, which are patterned alongside the venous sinuses, are also affected. Using a novel skull flat mounting technique, we show that the growth and expansion of meningeal lymphatics are perturbed in *Twist1* craniosynostosis models. Changes to the local meningeal environment, including hypoplastic dura and venous malformations, affect the ability of lymphatic networks to sprout and remodel. Dorsal networks along the transverse sinus are hypoplastic with reduced branching. By contrast, basal networks closer to the skull base are more variably affected, showing exuberant growth in some animals suggesting they are compensating for vessel loss in dorsal networks. Injecting molecular tracers into cerebrospinal fluid reveals significantly less drainage to the deep cervical lymph nodes, indicative of impaired lymphatic function. Collectively, our results show that meningeal lymphatic development is hindered in craniosynostosis, suggesting central nervous system waste clearance may be impeded.

## Introduction

The development and functions of meningeal lymphatic vessels (mLVs) have become subjects of intense interest because they are implicated in modulating the immunological state of the central nervous system (CNS) (Alves de Lima et al., 2020). These vessels reside in dura mater, the outermost meningeal layer encasing the CNS, along both the dural venous sinuses and meningeal arteries (Antila et al., 2017). Though the putative functions of para-arterial mLVs have yet to be discerned, perisinusoidal lymphatics along the venous sinuses have been found to uptake cerebrospinal fluid (CSF) (Louveau et al., 2018a, Ahn et al., 2019). During steady state, mLVs act in conjunction with the glymphatic system to transport macromolecules (Louveau et al. 2015), toxic brain metabolites (Xie et al. 2013, Lundgaard et al. 2016), amyloid-beta plaques (Da Mesquita et al., 2018), and dendritic cells (Louveau et al. 2018a) from the cranium to the deep cervical lymph nodes (dcLNs). As such, mLVs are essential for CNS homeostasis and normal brain functions, and their dysfunction is associated with neurodegeneration, cognitive impairment, and changes to neuroinflammatory processes (Da Mesquita et al., 2018; Louveau et al., 2018a; Alves de Lima et al., 2020).

Given their unique anatomical location, mLVs evaded our attention for centuries until their “rediscovery” in 2015 (Aspelund et al., 2015; Louveau et al., 2015). mLV networks have now been reported in humans, non-human primates, and rodents (Absinta et al., 2017). In mice, mLVs have a defined postnatal developmental pattern that involves endothelial cell migration, tube formation, and sprouting. mLVs start their growth by penetrating the dura through foramina at the base of the skull and begin forming preliminary networks along both the meningeal arteries and dural venous sinuses (Antila et al., 2017). As these networks mature, they continue to extend in a basal to dorsal direction. Para-arterial mLVs begin at the pterygopalatine artery and complete their growth near the top of the middle meningeal artery (MMA) by P28. Meanwhile, perisinus lymphatic networks begin developing along both the sigmoid (SgS) and petrosquamosal sinuses (PSS), before ascending the transverse sinus (TVS). Their development is completed by approximately P30, by which time coverage along the superior sagittal sinus (SSS) is complete (Antila et al., 2017; Izen et al., 2018).

These vessels share many commonalities with peripheral lymphatics, including the expression of Prox1, Lyve-1, and Podoplanin. In addition, mLVs also rely on Vegf-c/Vegfr3 signaling for their growth and maintenance (Antila et al., 2017). Vascular smooth muscle is postulated to be a local source of Vegf-c and the growth of mLVs along blood vessels is concurrent with venous smooth muscle development (Antila et al., 2017). By contrast, differential gene expression in meningeal versus peripheral lymphatic vessels has revealed that mLVs are unique according to gene pathways related to extracellular matrix (ECM) interactions, focal adhesion, angiogenesis, and responses to endogenous and exogenous stimuli (Da Mesquita et al., 2018). Thus, processes that govern the growth and maturation of mLVs are likely specific to their unique meningeal environment. Although the detailed mechanisms of mLV development remain largely unknown, these likely include tissue-specific interactions with blood vessels and dura. Utilizing animal models that possess structural changes to dura mater and/or the resident venous vasculature should help facilitate the discovery of these processes and their specific regulation of mLVs.

We recently reported that humans with craniosynostosis caused by mutations in the transcription factor *TWIST1* (Saethre-Chotzen syndrome) have dural venous sinus malformations, including loss and/or hypoplasia of the TVS, SgS, and jugular veins (Tischfield et al., 2017). Modeling the disorder in mice by inactivating *Twist1* in the cranial sutures and dura via *Sm22a-Cre* (*Twist1*^*Flx/Flx*^*:Sm22a-Cre*) unexpectedly revealed that *Twist1* regulates the growth and remodeling of the dural venous sinuses in a non-cell autonomous manner. It does so by promoting the development of osteoprogenitor cells and dura from cranial suture-derived mesenchymal progenitor cells, and these tissues produce growth factors (i.e. Bmp2/Bmp4) that act on the underlying venous endothelium (Tischfield et al., 2017). In addition to skull and vascular malformations, we also reported that dura was hypoplastic in affected embryos, but our analyses were limited to early embryonic stages preceding bone mineralization and skull development.

In the present study, we leverage *Twist1* craniosynostosis models that possess venous malformations and structural changes to dura to gain insight into processes that regulate the growth and remodeling of mLV networks. Using a novel skull flat mounting technique, we show that the growth and expansion of mLV networks is perturbed in craniosynostosis. Vessels that grow along the TVS show reduced branching and loss of network complexity. By contrast, lymphatic networks that grow along the SgS and PSS near the skull base are less affected and, in some animals, are hyperplastic with long vessels and numerous sprouts. Additionally, infusing tracer compounds into CSF reveals less drainage to the dcLNs, suggesting meningeal lymphatic networks are functionally compromised in craniosynostosis. Thus, mouse models for craniosynostosis can provide unique insights into processes that shape proper mLV development and function. Our findings now suggest that individuals with craniosynostosis may be at risk for impairments to meningeal lymphatic vessels and CNS waste clearance.

## Results and Discussion

We generated craniosynostosis models by inactivating both copies of *Twist1* in cranial mesenchyme, as previously reported (Tischfield et al., 2017). *Sm22a-Cre* is expressed by e10.5 in neural crest and cranial mesoderm, giving rise to dura and arachnoid tissue (El-Bizri et al., 2008, Tischfield et al., 2017). Lineage labeling also shows strong Cre activity in sutural mesenchyme, especially the coronal and sagittal sutures (Fig. 1A). These regions of non-ossified tissue separate the paired frontal and parietal bones, give rise to osteoblasts, and are commonly affected in Saethre-Chotzen syndrome (Twigg and Wilkie, 2015; Ottelander et al., 2021). To analyze skull malformations in higher detail, the skulls were imaged using three-dimensional x-ray microscopy. This revealed varying degrees of suture fusion coupled with calvarial bone loss in most animals (Fig. 1B), and skull length from the nasal to occipital bones was reduced by ∼25% (Fig. S1). In a small subset (13%, 4/30), the skull was intact and the animals had bilateral fusion of the coronal sutures and small dome shaped skulls. In ∼60% of animals (17/30), the coronal sutures showed bilateral fusion and bone loss was present along the dorsal midline where the sagittal suture is normally present, affecting the frontal, parietal, and interparietal bones (Fig. 1B). Areas that lacked proper mineralized bone instead contained a hardened, thin translucent material that lacked blood vessels but covered the underlying soft tissue. Also, focal bone loss was often observed in the region where the coronal suture is normally located (Fig. 2B). In the remaining animals, bone loss was more severe and stretched further away from the midline, and holes in the skull were sometimes present (Fig. S1). Thus, loss of *Twist1* in sutural mesenchyme via *Sm22a-Cre* models a severe form of craniosynostosis marked by premature fusion and/or partial agenesis of the cranial sutures, which can lead to regionalized bone loss.

**Figure 1:**
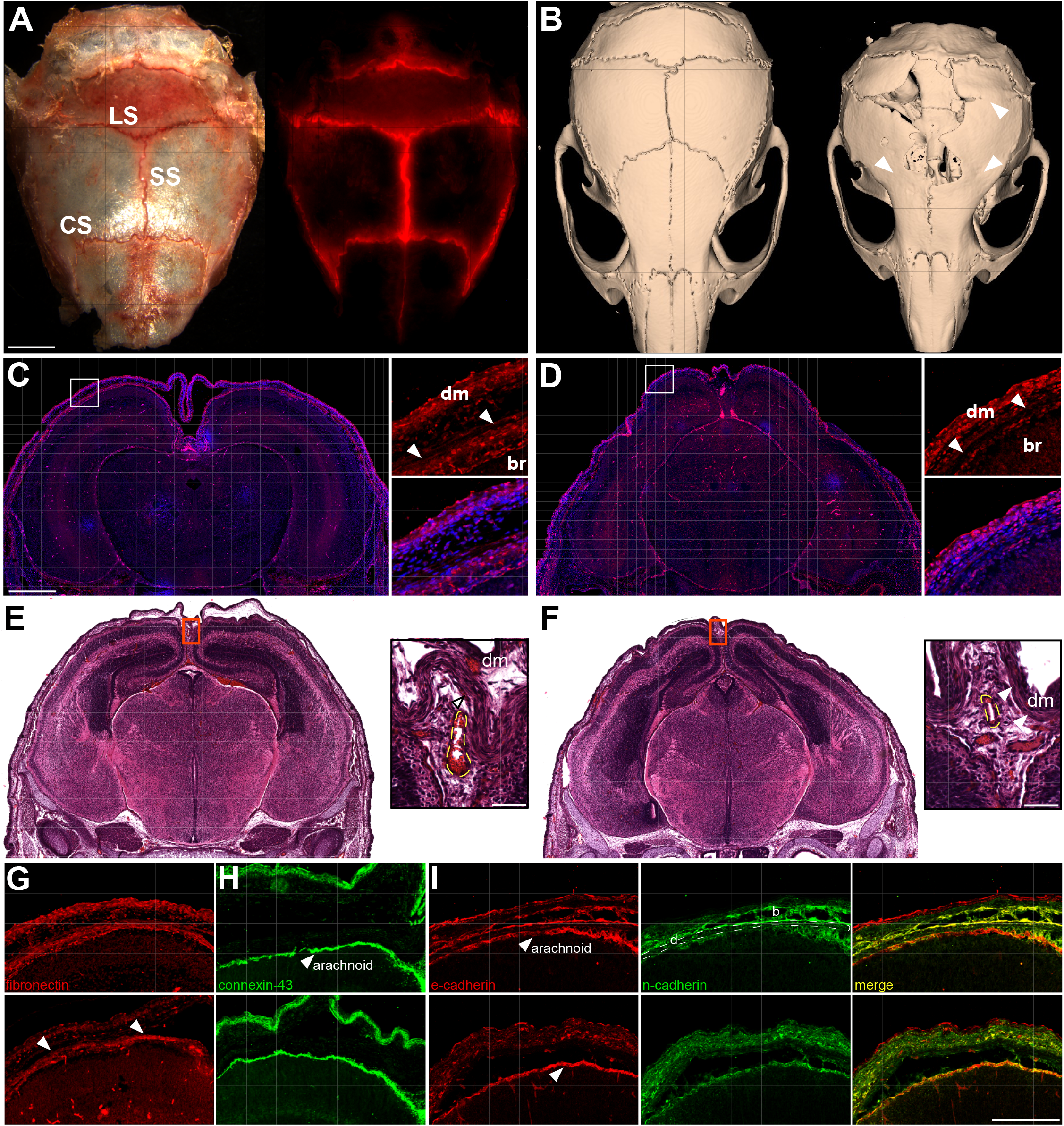
*Twist1*^*CS*^ animals have craniosynostosis and hypoplastic meninges. **(A)** *Sm22a-Cre* is active in sutural mesenchyme, as detected by the presence of red fluorescent protein (*Rosa26:Ai14*^*tdTomato*^). **(B)** Representative skull phenotype from a moderately affected *Twist1*^*CS*^ animal. Arrowheads denote fusion of the coronal and lambdoid sutures. **(C and D)** Coronal section through the heads of an e16.5 control **(C)** and severely affected *Twist1*^*CS*^ embryo **(D)**. Fibronectin (red) marks the meningeal extracellular matrix, which is markedly hypoplastic in severely affected embryos. **(E and F)** Coronal sections through the head of an e16.5 control **(E)** and a mildly affected *Twist1*^*CS*^ embryo **(F)** stained with hematoxylin and eosin. The meninges are preserved but hypoplastic periosteal dura and condensed osteogenic mesenchyme are observed at the dorsal midline and not well separated from the overlying dermis (dm, arrowheads). Dashed lines denote SSS hypoplasia. **(G-I)** e16.5 coronal sections adjacent to the dorsal midline from a control (top row) and mildly affected *Twist1*^*CS*^ embryo (bottom row). **(G)** The dura extracellular matrix is hypoplastic according to fibronectin staining (arrowheads). Connexin-43 **(H)** and e-cadherin **(I)** staining show the arachnoid membrane is intact. Bone (b) development, marked by n- and e-cadherin, is delayed and/or absent. SS=sagittal suture, CS=coronal suture, LS=lambdoid suture, d=dura, br=brain. Scale bars **(A)** 2.5mm, **(C)** 1mm, **(E)** 40µm.

**Figure 2:**
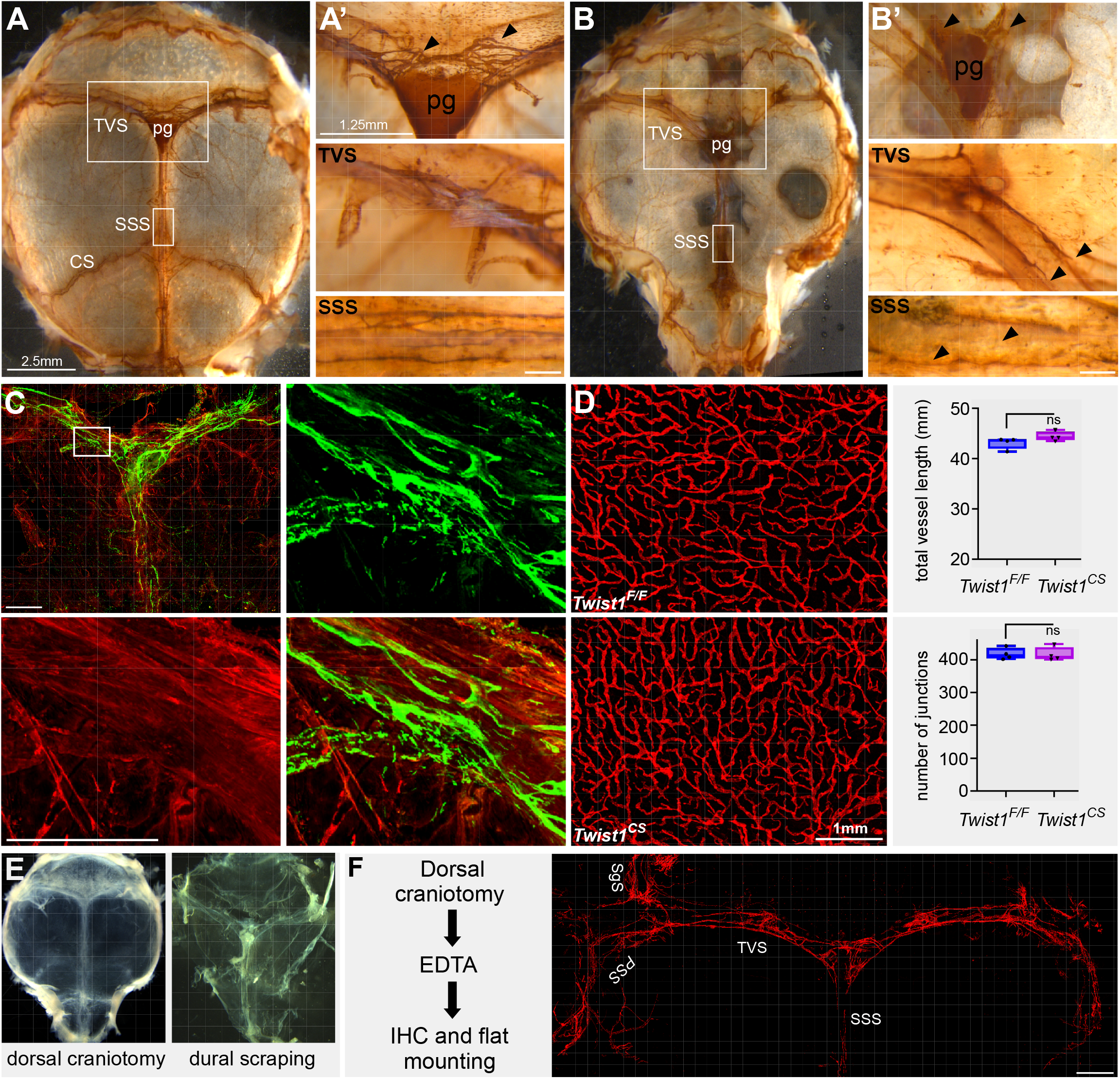
Dorsal meningeal lymphatic networks are affected in *Twist1*^*CS*^ animals. Dorsal craniotomies from a P60 control **(A)** and moderately affected *Twist1*^*CS*^ animal **(B)**. mLVs are depicted by chromogenic Lyve-1 staining (brown). Branched networks are present at the sinus confluence in controls **(A’**, arrowheads), and are missing in *Twist1*^*CS*^ animals **(B’**, arrowheads). Vessels along the TVS are hypoplastic in *Twist1*^*CS*^ and do not extend to the sinus confluence (arrowheads, middle panel), and vessel loss is noted along the SSS (arrowheads, bottom panel). **(C)** Dural scraping from a control animal expressing *Sm22a-Cre* and the *Rosa26:Ai14*^*tdTomato*^ reporter. *Sm22a-Cre* activity is detected in dura and vascular smooth muscle, but is absent in Lyve-1 positive mLVs (green). **(D)** Lymphatic networks in ear skin are normal in *Twist1*^*CS*^. **(E)** Traditional method for visualizing mLVs by scraping the dural membrane from the skull. **(F)** Representative example of the dorsal skull flat mounting method. The *Prox1*^*tdTomato*^ reported allele marks mLVs in red. TVS=transverse sinus, SSS=superior sagittal sinus, SgS=sigmoid sinus, PSS=petrosquamosal sinus, pg=pineal gland, CS=coronal suture. Scale bars, **(C)** 500µm **(F)** 1mm. ** *p<0*.*001*, *** *p<0*.*0001*, student’s t-test.

Periosteal dura is a thin portion of the dural membrane that directly underlies developing calvarial bones and sutural mesenchyme, and it secretes growth factors that are critical for skull development and suture maintenance (Ito et al., 2003; Lee et al., 2006). The dural venous sinuses and meningeal lymphatic vessels also reside in periosteal dura. We therefore examined the development of dura at or adjacent to the dorsal midline in e16.5 embryos, by which time the meninges have differentiated and bone mineralization is commencing.

In typical animals, the differentiation of the meninges starts at the skull base around embryonic day 12.5 (e12.5) and proceeds until ∼e14.5, by which time the three layers (dura, arachnoid, and pia) are observed from the skull base to the apex. Similarly, osteoblasts are formed from intramembranous ossification of condensed mesenchyme adjacent to dura. This process starts at eye level near the base of the head at ∼e12.5 and proceeds to the apex by e14.5-e15 (Deckelbaum et al., 2012; DeSisto et al., 2020). In affected embryos, meningeal tissue closer to the skull base was relatively normal and three layers were recognized. In severely affected embryos, however, condensed mesenchyme and dura was largely absent along the dorsolateral margins of the head and dorsal midline, and the apical expansion of bone growth was halted. The dural extracellular matrix was missing as fibronectin staining was largely absent (Figs. 1C, D). In more common, mildly affected embryos, the dura and meninges were intact but became noticeably hypoplastic at or near the dorsal midline. Unlike controls in which condensed osteogenic mesenchyme was clearly separated from the overlying dermis, these embryos showed less periosteal dura and osteogenic mesenchyme without clear separation from the overlying dermis (Figures 1E, F). The superior sagittal sinus (SSS) was hypoplastic but still enveloped by dura, and all affected animals had segmental or complete unilateral loss of the TVS (Fig. S2), as reported previously in mice and humans (Tischfield et al., 2017). Arachnoid tissue was also present, as detected by the presence of e-cadherin and connexin-43, but fibronectin expression was reduced in regions where dura was hypoplastic, and n-cadherin positive osteoblasts had not yet developed (Figs. 1G-I). These results show that calvarial bone loss parallels the absence of dura, and loss of periosteal dura is more significant dorsolaterally and/or at the dorsal midline of the head where the dural venous sinuses and mLV networks reside.

### Meningeal lymphatic networks are affected in *Twist1* craniosynostosis models

Typically, mLVs are examined by carefully scraping dura off the skull, followed by immunohistochemistry, flat mounting, and imaging (Louveau et al., 2015). This was unfeasible in *Twist1*^*Flx/Flx*^*:Sm22a-Cre* (*Twist1*^*CS*^) animals since the hypoplastic dura did not stay intact during scraping. We instead removed the dorsal half of the skull, performed chromogenic immunohistochemistry to stain vessels with Lyve-1, and imaged the lymphatic networks under a stereomicroscope. In control animals, mLVs were patterned along the SSS and TVS as reported (Antila et al., 2017). The networks were more complex with numerous branches at the sinus confluence and at particular locations along the TVS (Fig. 2A, A’). These regions are proposed to be “hotspots” specialized for the uptake of CSF (Louveau et al., 2018a). By contrast, meningeal lymphatic networks were poorly developed in *Twist1*^*CS*^ animals (Figs. 2B, B’, n=3). Networks along the TVS appeared less complex and/or atrophic, and growth towards the sinus confluence was impeded. Vessels along the SSS were also hypoplastic and/or missing (Fig. 2B’). Lineage labeling showed that *Sm22a-Cre* is not expressed in mLVs (Fig. 2C). Furthermore, we previously reported that *Twist1* is not expressed in meningeal endothelial cells, and *Twist1*^*FLX/FLX*^*:Tie2-Cre* embryos have normal vascular development with no signs of edema (Tischfield et al., 2017). The development of lymphatic networks in other tissues, such as ear skin, was normal (Fig. 2D). Thus, the observed phenotypes are non-cell autonomous and are due to changes in the local environment that hinder the proper growth and expansion of lymphatic networks.

### The growth of meningeal lymphatic networks is perturbed in *Twist1*^*CS*^ animals

Overall, it was difficult to visualize vessels at the sinus confluence or along the SSS in *Twist1*^*CS*^ mice using chromogenic staining and a stereomicroscope. We therefore sought to develop alternative, high-resolution methods for imaging meningeal lymphatic networks (Fig. 2D, E, Fig. S3). To image dorsal networks along the TVS and SSS, we adapted a previous method that involved removing the dorsal half of the skull, above the temporal bone, with the meninges still attached (Louveau et al., 2018b). The skullcaps were post-fixed and decalcified to remove auto fluorescence, and the tissue was stained using free-floating immunohistochemistry. Incisions were made along the corners of the frontal and occipital bones and the tissue was flat mounted onto slides. This technique provides superior resolution compared with chromogenic immunohistochemistry (Fig. 2D) and allows the option for 3D reconstructions.

We used this high-resolution technique to determine if mLV development was affected in juvenile animals at postnatal (P) day 16 when mLVs are actively growing and remodeling along the TVS. The TVS was visualized using alpha-smooth muscle actin (aSMA) to mark the thin layer of smooth muscle that envelopes the venous endothelium, and mLVs were stained with Lyve-1. In controls, meningeal lymphatic networks exhibited numerous sprouts along the TVS (Fig. 3A). By contrast, networks in *Twist1*^*CS*^ animals mostly consisted of long, unbranched vessels that were devoid of sprouts, and coverage along the TVS was reduced (Fig. 3B, n=3). In severely affected adult animals, the lymphatic networks were markedly hypoplastic along the TVS and SSS and largely missing around portions proximal to the confluence. aSMA staining revealed that hypoplastic tissue surrounding the confluence and underlying regions of bone loss contained myofibroblasts, suggesting it was fibrotic. Tangled bundles of blood vessels were commonly observed and Lyve-1 positive macrophages were abundant. These macrophages normally reside in the leptomeninges (Mundt et al., 2019; Van Hove et al., 2019), and their presence suggests arachnoid tissue remained attached to the skull and thin layer of hypoplastic dura in more severely affected *Twist1*^*CS*^ animals. Venous smooth muscle coverage appeared patchy in severely affected animals compared to controls, and aSMA-positive cells were sometimes present in tissue surrounding the proximal portions of the TVS adjacent to the SgS and PSS, suggesting they failed to migrate (Figs. 3C, D). Interestingly, at the sinus confluence, hyperplastic lymphatic vessels were sometimes observed (Fig. 3D). Overall, these results indicate that the growth and sprouting of mLVs is affected in *Twist1*^*CS*^ animals, accounting for loss of meningeal lymphatic networks in adult animals.

**Figure 3:**
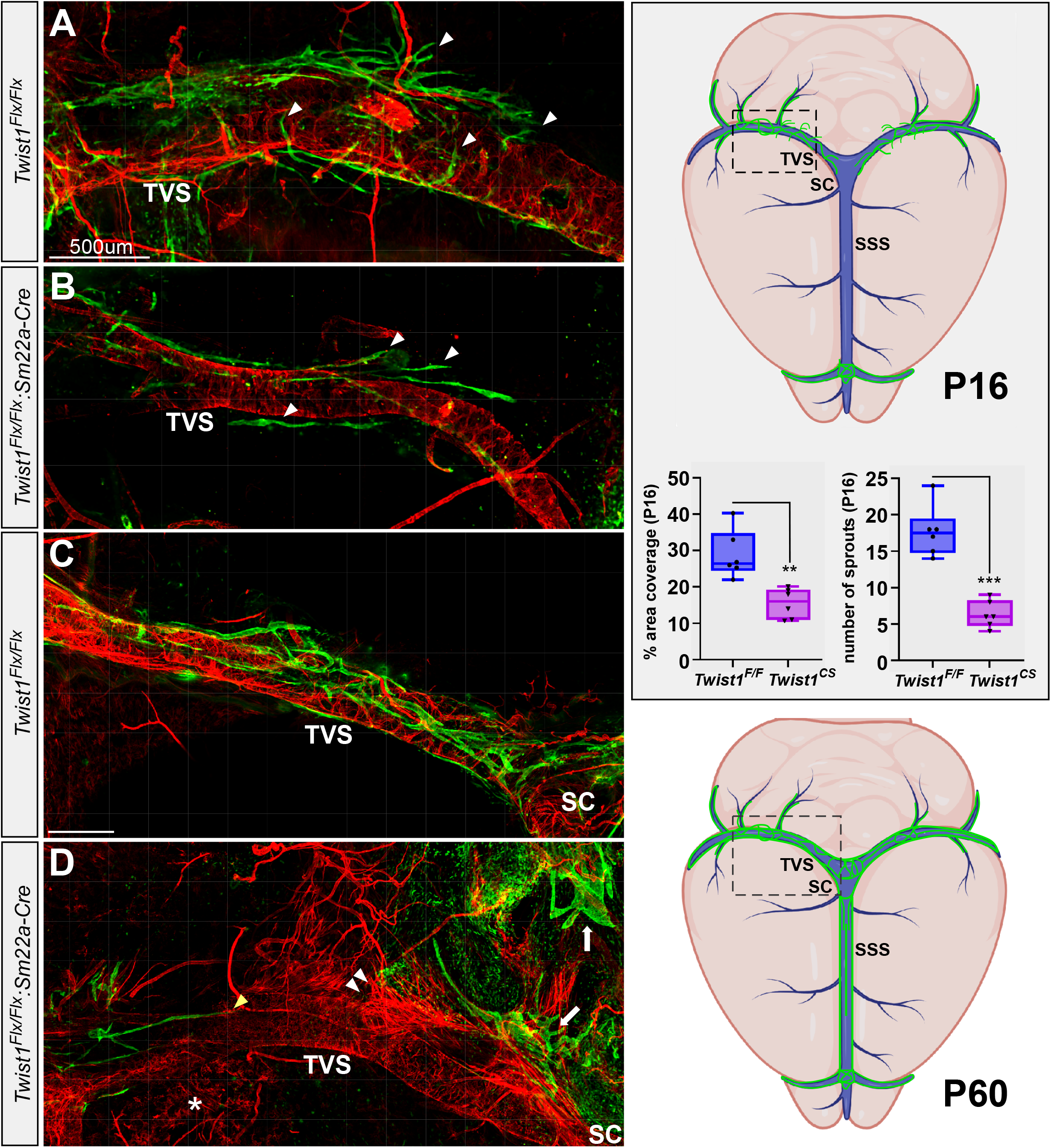
Growth and expansion of meningeal lymphatic networks is affected in *Twist1*^*CS*^. **(A and B)** Lyve-1 (green) denotes developing mLVs along the TVS (marked by α-SMA, red) in a P16 control **(A)** and moderately affected *Twist1*^*CS*^ animal **(B)**. Developing lymphatic networks show numerous sprouts along the TVS in controls, and significantly less in *Twist1*^*CS*^. Network expansion is reduced according to percent area coverage at P16 (n=3, each side of TVS). Right diagrams illustrate the development of meningeal lymphatic networks along the venous sinuses at P16, and mature networks at P60. **(C)** Mature lymphatic networks along the TVS in a P90 control and severely affected *Twist1*^*CS*^ animal **(D)**. mLVs are largely absent in the affected animal, and only a few vessels are present along the initial segments of the TVS (yellow arrowhead). Smooth muscle coverage along the TVS is patchy and α-SMA positive cells are seen in adjacent dura (asterisk). Ectopic blood vessels are present and α-SMA positive myofibroblasts surround the sinus confluence (SC). Hyperplastic lymphatic vessels are sometimes present at the sinus confluence (sc) in severely affected animals (white arrows) along with Lyve-1 positive macrophages.

### Basal meningeal lymphatic networks are variably affected in *Twist1*^*CS*^ animals

Although dorsal skull flat mounts provide excellent resolution of vessels along the SSS and TVS, basal networks present along the SgS and PSS were often damaged during the procedure. We therefore analyzed basal meningeal lymphatic networks using a variation of the aforementioned technique (Fig. S3). The skull was harvested and bisected along the dorsal midline, treated with the same post-fixation and decalcification conditions, and excess muscle, fat, hair and connective tissue were removed. The dural tentorium was cut off, the optic nerves were removed from their sheathes, and incisions were made on the corners of the rounded skull to permit flat mounting of the softened tissue onto a slide. The temporal bone was carefully removed in order to not disrupt basal lymphatic networks that reside along the SgS and PSS, which are located along the dorsal and lateral margins of this bone, respectively (Fig. S3). In some animals, it was necessary to thin the skull by peeling off the outermost layer of the calvarial bone.

These “basal preparations” fully preserve mLV networks along the SgS and PSS, and also allow imaging of mLVs along arteries near the skull base. Moreover, the vessels along the TVS are also preserved, but care must be taken to preserve networks at the lateral edges of the sinus confluence. Overall, both techniques provide superior resolution to dural flat mounting. They are not subject to tears and creases that can occur from handling the thin dural tissue that has been scraped from the skull, and perfectly preserve the 3D architecture of the lymphatic networks. For these analyses, we crossed *Twist1*^*CS*^ animals with a *Prox1*^*tdTomato*^ BAC-transgenic reporter line that provides bright red fluorescence in mLVs (Fig. 4A)(Hong et al., 2016).

**Figure 4:**
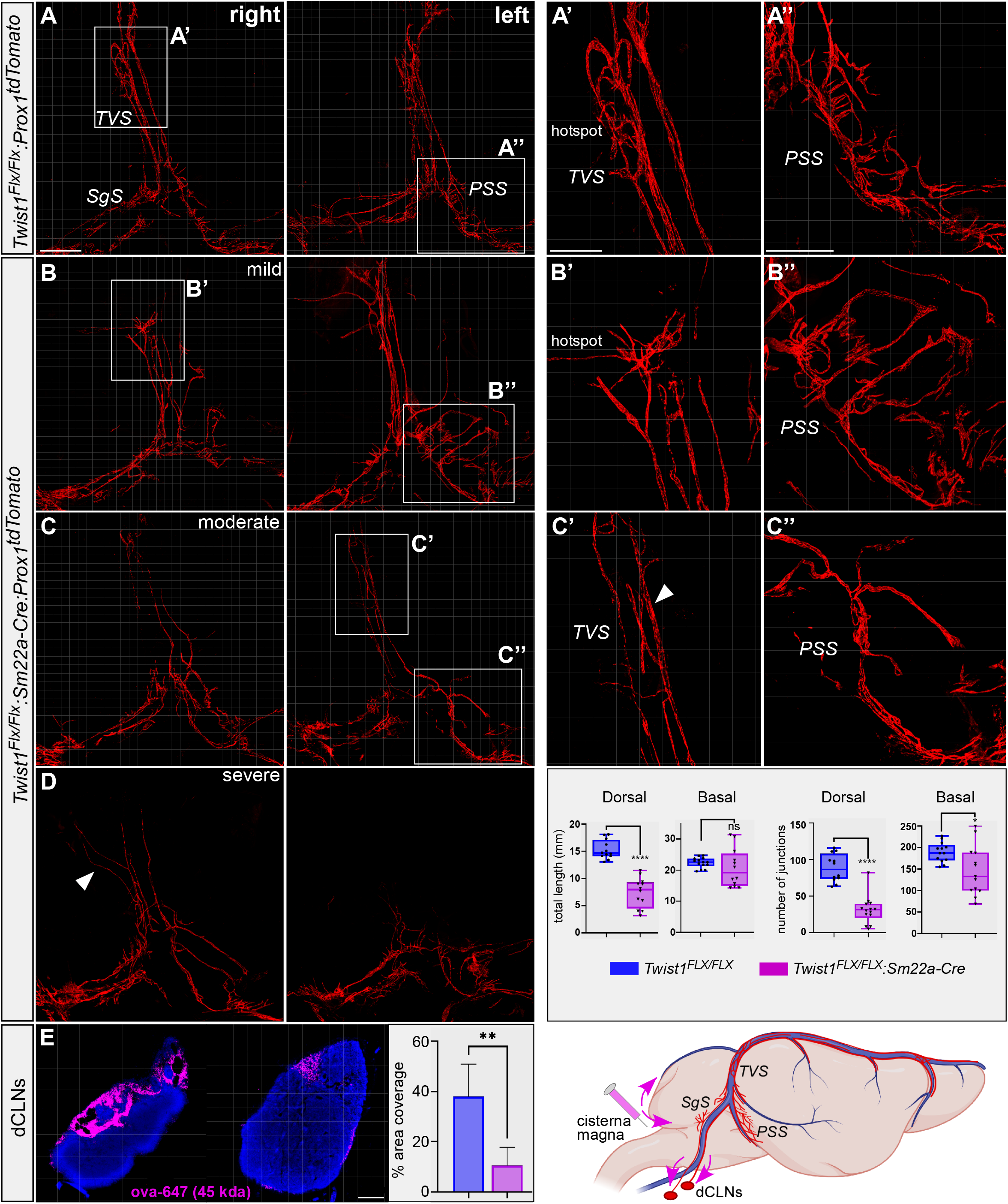
Basal lymphatic networks are variably affected in *Twist1*^*CS*^ animals. **(A)** Basal preparation depicting RFP labeled mLVs in a control *Twist1*^*FLX/FLX*^*:Prox1*^*tdTomato*^ adult. Boxed regions denote dorsal networks along the TVS and basal networks along the PSS. Left and right panels show mLVs present on each half of the bisected skull. Dorsal vessels along the TVS show extensive branching at “hotspot” regions **(A’)**. Basal networks along the PSS also show increased complexity as shown in right panel **(A’’). (B-D)** mLV networks in mildly affected **(B)**, moderately affected **(C)**, and severely affected **(D)** *Twist1*^*CS*^*:Prox1*^*tdTomato*^ animals. In mildly affected animals, networks along the TVS appear normal, but mLVs are missing on the side with segmental loss of the TVS **(B’)**. Basal lymphatic networks show excessively long vessels with numerous branches (**B’’**). Lymphatic networks along the TVS in moderately affected animals are hypoplastic and less complex (compare **A’** and **C’**) with loss of “hotspots”. As shown in **(C’’)**, basal networks are variably affected but typically less complex with fewer branches, and abnormally long vessels can be present. In severely affected animals, networks along the TVS are extremely hypoplastic and/or missing. Lymphatic vessels are present along ectopic veins (arrowhead in **D**). Basal networks show variable affection. In this animal, the left side shows long, unbranched vessels, whereas the network on the right is hyperplastic and disorganized. Quantifications for dorsal networks correspond to those vessels present in the boxed TVS region in panel A, and basal networks correspond to vessels present in boxed PSS regions, including those present along the SgS. (E) The amount of tracer is reduced in *Twist1*^*CS*^ dcLNs (right) following infusions via the cisterna magna. Scale bar= 1mm (A) and 200um (E).

In mildly affected animals, growth along the TVS was more comparable to control littermates as branched networks were observed at “hotspot” regions; however, vessel growth was still absent in areas where the TVS was missing (Fig. 4B, B’). Interestingly, basal networks along the PSS were exuberant in these mildly affected animals, containing long vessels with numerous branches (Fig. 4B n=2). In moderately affected animals, hypoplastic networks with rudimentary “hotspots” and less branching were present along the intact TVS, but were largely absent and/or atrophic on the side where the TVS was missing (Fig. 4C, C’, n=3). Basal lymphatic networks along the PSS were less affected than dorsal networks. Long vessels were observed but branching and complexity was reduced compared to more mildly affected animals and controls. In severely affected animals (n=2), mLV growth and complexity along the TVS was significantly affected, and few if any sprouts and branches were observed. mLVs were also present along ectopic veins that branched from the remaining segments of the TVS (Fig. 4D). Basal networks along the PSS were variably affected. Similar to mild/moderately affected animals, long unbranched vessels were sometimes observed, and disorganized hyperplastic networks could accompany these on the contralateral side (Fig. 4D). In general, animals that were missing dorsal networks had exuberant basal networks, suggesting they were compensating for vessel loss along the TVS. The growth of mLVs along the sigmoid sinus was also more robust in severely affected *Twist1*^*CS*^ animals versus controls (Fig. 4D). Finally, vessels along the PPA and MMA were mildly affected; coverage along distal, dorsal portions of the MMA towards to skull apex appeared reduced in regions where dura was hypoplastic (Fig. S4).

### Lymphatic drainage to the deep cervical lymph nodes is reduced in *Twist1*^*CS*^ animals

Meningeal lymphatics drain macromolecules and waste from CSF, and facilitate the trafficking of immune cells to the dcLNs (Louveau et al., 2018a). Although the main collection point of meningeal lymphatic drainage is the dcLNs, these vessels also drain to the superficial cervical lymph nodes (scLNs), albeit to a lesser extent (Aspelund et al., 2015). To determine if drainage to the dcLNs was affected, we infused a 45kDa Ovalbumin-647 tracer into the CSF by accessing the subarachnoid space through the cisterna magna. We selected mild/moderately affected animals in which the meninges and subarachnoid space were preserved. The amount of tracer in the dCLNs was significantly reduced in *Twist1*^*CS*^ animals (Fig. 4E), whereas the amount of tracer in the scLNs was comparable (data not shown). In addition, the dcLNs showed unilateral hypoplasia in some animals, suggesting lymphatic drainage was ipsilateral or passaged through alternative routes. These results suggest that both the growth and functions of meningeal lymphatic networks are compromised in craniosynostosis.

Our results show that changes to the local meningeal environment are sufficient to disrupt the growth and remodeling of mLVs in craniosynostosis. The model we have chosen for our current analysis is overall more affected than what is typically observed in humans, and shows similarities with Sweeney-Cox syndrome, a severe craniofacial disorder caused by dominant-negative mutations in *TWIST1* (Kim et al., 2017; Takenouchi et al., 2018). However, homozygous *Twist1*^*FLX/FLX*^*:Sm22a-Cre* animals model venous malformations that are absent in heterozygous *Twist1*^*FLX/WT*^*:Sm22a-Cre* animals, the latter of which more closely mimic skull phenotypes found in humans (Tischfield et al., 2017). Furthermore, more mildly affected animals with bilateral coronal suture fusion approximate the human condition and *Twist1*^*FLX/WT*^*:Sm22a-Cre* heterozygous animals. Thus, the lymphatic phenotypes we describe here are likely to be present in humans to varying degrees, especially those who have more severe forms of craniosynostosis with loss or hypoplasia of the dural venous sinuses. Work is underway to investigate meningeal lymphatic phenotypes in heterozygous *Twist1*^*FLX/WT*^*:Sm22a-Cre* animals, and other forms of craniosynostosis caused by activating gene mutations in *FGFR2* (Johnson and Wilkie, 2011).

The growth of mLVs is dependent upon Vegf-c, which is expressed by venous and arterial smooth muscle in the meninges (Antila et al., 2017). In agreement, mLVs were rarely observed in regions where the TVS was missing in affected animals, whereas vessel growth was observed along ectopic veins that developed in the absence of proper venous sinuses. This suggests smooth muscle derived growth factor signaling from these vessels was sufficient to induce lymphatic growth. Smooth muscle coverage appeared normal on the middle meningeal arteries (which are preserved in these animals), and we did not observe overt changes to meningeal lymphatics that grew alongside these vessels, with the exception of the more dorsal portions of the MMA where dura was hypoplastic. Thus, some of the observed changes to lymphatic networks may be attributed to venous malformations and attenuated Vegf-c signaling. In addition, hypoplastic dura and loss of the extracellular matrix is likely to affect the development of lymphatic networks. Notably, the development of dermal lymphatic networks in mid-gestation embryos relies on extracellular activation of integrin-β1, which can bind and activate Vegfr3 independent from Vegf-c (Planas-Paz et al., 2012). Also, the migratory abilities of dermal lymphatics in mid-gestation embryos are influenced by mechanical forces regulated by tissue stiffness (Frye et al., 2018), which may be altered in more severely affected animals with hypoplastic, fibrotic dura. Thus, mLV phenotypes in *Twist1*^*FLX/FLX*^*:Sm22a-Cre* animals may manifest from combinatorial changes to the surrounding environment that impinge upon Vegfr3 activation, and the growth and sprouting of lymphatic networks. Future studies will examine changes to these processes in our models.

In even mildly/moderately affected *Twist1*^*FLX/FLX*^*:Sm22a-Cre* animals, lymphatic drainage to the dcLNs was significantly diminished. This can be expected to affect immune cell trafficking as well as CNS waste clearance, especially because ablating mLVs affects the functions of the brain’s perivascular waste clearance system (i.e. the glymphatic system) (Da Mesquita et al., 2018; Louveau et al., 2017). Impaired CNS waste clearance is associated with the accumulation of amyloid-beta plaques and cognitive impairment (Iliff et al., 2012; Da Mesquita et al., 2018), whereas altered meningeal immune cell trafficking can also affect brain function and behavior in addition to neuroinflammatory processes (Louveau et al., 2018a; Alves de Lima et al., 2020). In aging animals, meningeal lymphatic networks also naturally deteriorate; dorsal vessels regress whereas basal vessels become hyperplastic (Ahn et al., 2019). Interestingly, hyperplastic basal networks were seen in a subset of animals in the present study. Notably, it is unknown if craniosynostosis may be associated with a higher prevalence of Alzheimer’s disease or other forms of neurodegeneration that lead to cognitive decline and dementia. Likewise, to our knowledge, changes to neuroinflammatory processes have not been reported or studied in craniosynostosis. Given that craniosynostosis is associated with venous hypertension and impaired CSF drainage, our new findings imply that many individuals with craniosynostosis, especially syndromic forms, may unknowingly be at heightened risk for cognitive decline due to cerebrovascular malformations, impaired waste clearance, and/or changes to neuroinflammatory processes resulting from altered immune surveillance.

In the present study, we have developed new methods to facilitate the characterization of meningeal lymphangiogenesis. Although dural scrapings provide a fast and cost-effective method for visualizing mLVs, this method may not be suitable for all models, especially those possessing meningeal malformations. Our innovative skull flat mounting technique is a superior alternative, as it perfectly preserves meningeal lymphatic networks and provides high resolution imaging in their native environment. Moreover, it does not suffer from imaging artifacts that arise from rips, creases, and folding which may occur from scraping dura from the skull and/or flat mounting the tissue onto slides. However, it does require more time to fix, soften, and stain the tissue for imaging. Thus, the best techniques for studying mLVs may depend upon the specific animal model(s) and the types of questions that need to be addressed.

---

**Supplemental Figure 1: Skull development in a severely affected *Twist1***^***CS***^ **animal**. Loss of mineralized bone is more extensive in severely affected *Twist1*^*CS*^ animals. In this animal (P60), large gaps of non-ossified tissue are found in regions where the coronal (cs), frontal (fs), and sagittal sutures (ss) normally develop. Meningeal hypoplasia is also more severe in these animals, as seen in figure 1d. The length of the skull, as measured from the frontal bone to the occipital bone (dashed line), is reduced by approximately 25% in *Twist1*^*C S*^ animals. *** *p<0*.*0001*, students t-test.

**Supplemental Figure 2**: ***Twist1***^***CS***^ **animals have hypoplastic dural venous sinuses with segmental loss of the TVS:** Dorsal skull flat mounts from a P16 control (left) and *Twist1*^*CS*^ animal (right). The dural venous sinuses and mLVs are visualized according to alpha-smooth muscle actin (red) and Lyve-1 (green) staining, respectively. All *Twist1*^*CS*^ animals have unilateral or segmental loss of the TVS (arrowheads). The intact TVS is typically hypoplastic and tortuous and the SSS is hypoplastic in all animals. Lyve-1 macrophages surround the sinus confluence in moderate/severely affected animals. Note the absence of lymphatic vessel sprouting along the intact TVS and contralateral remaining segment in the *Twist1*^*CS*^ animal compared to the control.

**Supplemental Figure 3: Workflow and procedure for preparing basal skull flat mounted tissue: (1)** Following immunohistochemistry, carefully gross the basal sample by cutting off the nasal mucosa, maxilla, and mandible. Next, remove the cartilage, muscle, fat, and excess connective tissue on the outer surface of the skull. If done well, you should receive results similar to (2). Make sure to clean the outer cranial surface with care so to not tear or puncture decalcified skull bone, as this will damage the meninges inside of the skull. **(2)** Use micro-scissors to cut off the tentorium and remove the optic nerve sheath inside the cranium. Perform this procedure under a fluorescent microscope. Switch between bright field and epifluorescence to plan your cuts and prevent accidental damage to the mLVs. On the outside of the cranium, cut or peel off the zygomatic arch. Results should approximate (3). **(3)** Inspect work and adjust cuts/cleaning as necessary. **(4)** While switching between fluorescence and bright-field, cut out areas of the occipital (ob), parietal (pb), and frontal bones (fb) without disturbing the mLVs, as indicted by the red dashed lines. **(5)** Use micro-scissors and extreme care to remove the inner ear (ie) bone. Switching between fluorescence and bright-field will help prevent damage to mLVs. **(6)** Inspect work and adjust cuts/cleaning as necessary. **(7)** Place tissue on a slide with the meningeal side (inner cranium) facing up, and create a large covering of mounting media around the sample. This can be done by withdrawing 600µl of mounting media. Pipet 200-300µl on the slide and overlay the sample. Then place 300-400µl of mounting media on top of the sample. **(8)** Apply coverslip, allow to dry, and image. Samples can be refrigerated and stored long term.

**Supplemental Figure 4: mLV coverage along the middle meningeal artery is mildly reduced in *Twist1***^***CS***^: *Prox1*^*tdTomato*^ labeled mLVs along the pterygopalatine (PPA) and middle meningeal arteries (MMA) in a P60 control (left) and *Twist1*^*CS*^ animal (right). Coverage along dorsal segments of the MMA towards the skull apex is patchy and reduced in *Twist1*^*CS*^ animals (arrowheads). Otherwise, arterial mLV coverage is relatively normal in *Twist1*^*CS*^ animals.

## Materials and Methods

### Animals

The following transgenic mice were used: *Twist1*^*FLX*^ (RRID:MMRRC_016842-UNC), *Prox1*^*tdTomato*^ (RRID:MMRRC_036531-UCD), *Rosa26:Ai14*^*tdTomato*^ (RRID:IMSR_JAX:007914). For all experiments, male and female mice were included. Animals were maintained on a mixed genetic background (C57Bl/6;FVB;CD1). Embryos obtained from timed matings were considered 0.5 days old upon observance of a plug. Experiments were approved and carried out under IACUC protocol PROTO201702623 (MAT).

### Antibodies

The following antibodies were used: Lyve-1 (1:300, Abcam, ab14917, RRID:AB_301509), RFP (1:1500, Rockland, 600-401-379, RRID:AB_2209751), Fibronectin (1:250, Abcam, ab2413, RRID:AB_2262874), αSMA (1:300, Sigma, C6198, RRID:AB_476856). Nuclei were visualized using Hoechst staining (1:2000, Thermofisher H3570).

### Immunohistochemistry

Heads from E16.5 embryos were decapitated and fixed overnight in 4% PFA at 4°C. The heads were then washed and dehydrated in steps of 50% and 80% ethanol prior to paraffin embedding. 10µm paraffin sections were collected for staining and images were obtained using a LSM700 confocal microscope with a 20× 0.8 NA objective. For chromogenic 3,3’-diaminobenzidine (DAB) staining, the SignalStain DAB substrate kit was used (Cell Signaling Cat# 8059) according to manufacturer’s instructions. Briefly, following application of the primary antibody (Lyve-1), 800µl of ant-rabbit horseradish peroxidase was added to the excised dorsal skull tissue for 30 minutes in a 24 well plate at room temperature. 30µl of DAB was added to 1ml of the DAB substrate solution. The solution was added to the tissue and allowed to incubate for 2-4 minutes until the signal was detectable but not saturated. The solution was then discarded and the tissue was washed in distilled water.

### Dorsal and basal skull flat mount preparations

The animals were perfused with 4% paraformaldehyde, heads decapitated, and all skin and musculature were removed from the dorsal half of the skull. The lower jaw was removed by inserting scissors into the oral cavity to cut the mandible and attached muscles. Angled micro scissors were used to cut laterally from the cisterna magna along the temporal bone to each orbit, and the nasal bones were severed to isolate the calvarium. The tissue was post-fixed in 2% PFA overnight at 4° Celsius, washed in PBS, and then treated with Dent’s Fix (80% methanol, 20% DMSO) overnight at 4° Celsius. Skulls were next decalcified in 14% EDTA for seven days at 4° Celsius and then cleaned of fat, muscle, and connective tissue. Afterwards, the skulls were incubated with primary antibodies for four days, followed by secondary antibodies for two days at 4° Celsius in 0.5% triton with 10% normal goat serum. Incisions were made at the corners of the frontal and occipital bones, and the tissue was flat mounted onto slides in mounting medium (SouthernBiotech Fluoromount-G) and cover slipped. Basal flat mounts, which are described in detail in supplementary figure 3, were otherwise similar to the dorsal flat mount preparation. The skull was bisected along the dorsal midline into halves by making an incision along the occipital bone midline, inserting the tips of scissors into the nasal bones, and opening the scissors to split the skull.

### Skull 3D X-ray microscopy (computed tomography)

Animals were sacrificed via transcardial perfusion with 4% paraformaldehyde (PFA). The heads were then decapitated and post-fixed overnight in 4% PFA. Hair and skin were removed prior to imaging. Images were obtained using a Bruker Skyscan 1272 X-ray microscope. The following scan conditions were used: Image pixel size=13.5um, camera=1632 columns x 1092 rows, rotation step=0.4 degrees, Frame averaging=3, Filter=1mm Al. The resulting images were reconstructed and converted to dicom format with Skyscan Ctan software. Dicom files were opened in Vivoquant for segmentation of teeth and bone from less dense soft tissues.

### Microscopy and lymphatic vessel quantifications

Images were acquired on a LSM700 confocal microscope using a 10× 0.3 NA objective. For dorsal and basal lymphatic preparations in figures 3 and 4, maximum intensity projections (MIPs) were obtained using 7µm z-stacks. For the quantification of sprouts along the TVS, these were manually counted from three control and experimental animals. Values were obtained from vessels present along the left and right transverse sinuses. Each value was imputed separately for statistical analysis (n=3 animals, six data points). The total vessel length and number of junctions for both the dorsal and basal networks in figure 4 were analyzed from MIPs using the AngioTool plug-in for ImageJ. For dorsal networks, a region of interest was centered over the transverse sinus, extending to the sinus confluence. For basal networks, a region of interest was centered over vessels along both the sigmoid and petrosquamosal sinuses (refer to figure 4a). The following settings were used in Angiotool: Dorsal networks, value diameter and intensity=12 and for basal networks, value diameter and intensity=9, fill small holes=80. Values were obtained from at least one half of the skull for vessels comprising dorsal and basal networks from six control and seven experimental animals.

### Infusion of molecular tracers into cerebrospinal fluid

Animals were anesthetized using ketamine/xylazine (100mg/kg). An incision was made on the midline of the head at the occipital crest. Once the skin was excised, curved forceps were used to break through the superficial connective tissue to reveal the underlying muscle. The muscle was carefully separated along the midline to expose the opening to the cisterna magna, and care was taken to not induce tears or bleeding. A 45kDa ovalbumin-647 tracer (Molecular Probes, Invitrogen) was mixed in artificial cerebrospinal fluid (20µg/ml solution), and 5µl of solution was loaded into a 10µl Hamilton syringe attached to polyethylene tubing and injected into the cerebrospinal fluid via the cisterna magna. The solution was injected using a 28-gauge needle and a Nanomite infusion system (Harvard Apparatus) with an injection rate of 2.5µl/min. Animals were kept on supplemental oxygen throughout the duration of the experiment to stabilize breathing and minimize hypercapnia, and the needle was glued in place to prevent depressurization. After 30 minutes had passed, the needle was removed and the animals were immediately euthanized by transcardial perfusion.

### Cervical lymph node imaging and quantifications

Animals were perfused with 4% PFA and the superficial and deep cervical lymph nodes were dissected under fluorescence. The tissue was allowed to post-fix for 12 hours in 2% PFA, prior to sinking in 30% sucrose and embedding in Neg-50 medium. 20µm sections were cut from three control and three experimental animals. The sections were imaged using a Leica M165FC stereomicroscope equipped with a 1x objective and DFC7000T camera. Ten 20µm sections per animal from the deep cervical lymph nodes were thresholded and analyzed by tracing out the tissue sections and calculating the percent area coverage. The values were averaged for each animal to obtain a single value representing the average percent area coverage for the deep cervical lymph nodes. Representative images were selected and imaged using a LSM800 confocal microscope with a 20× 0.80 NA objective.

### Statistics

Statistics were performed using GraphPad Prism 9.0. For all analyses, unpaired student t tests with Welch’s correction were performed.

## Author Contributions

Conceived and designed project (PA and MAT). Performed experiments and analysed data (PA, MM, MAT). Wrote and edited manuscript (PA, MM, MAT).

## Acknowledgments

The authors would like to thank Young Kwon-Hong (University of Southern California) for providing *Prox1*^*tdTomato*^ mice and the Rutgers Molecular Imaging Center (D. Adler and P. Buckendahl) for assistance with skull 3D x-ray microscopy. Michael Falen and Kush Desai for assistance with mouse dissections and immunohistochemistry. VE Abraira for providing images from BioRender used in figure 3. Marianne Polunas and the Rutgers Research Pathology Services Core for assistance with embryo embedding and hematoxylin/eosin slide preparation.

