## Supplementary figures and images for "The growth and expansion of meningeal lymphatic networks are affected in craniosynostosis"

### Supplemental Figure 1

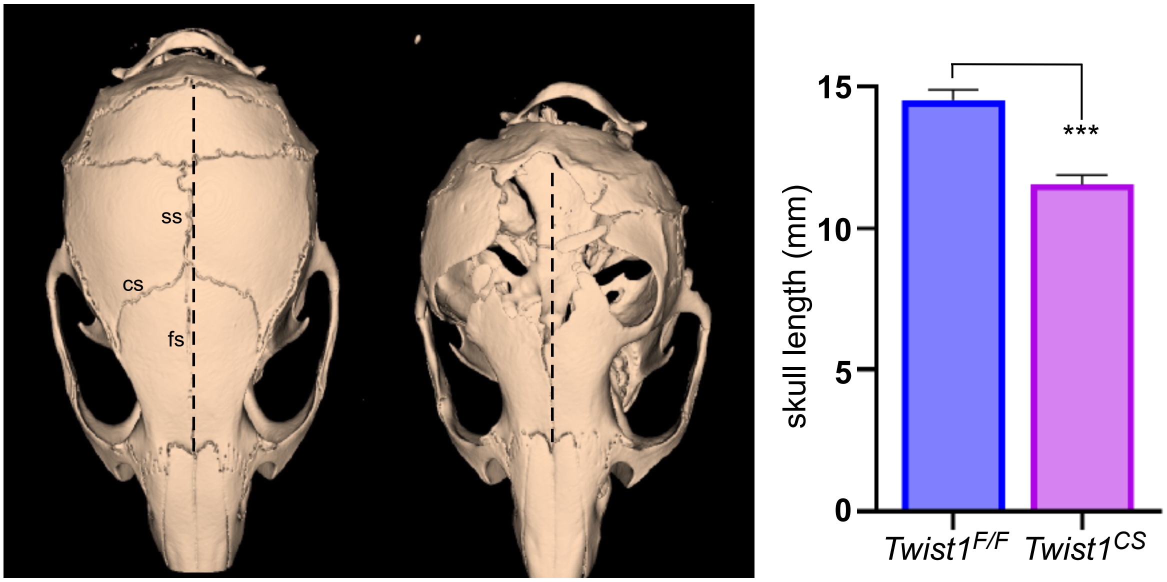

### Supplemental Figure 2

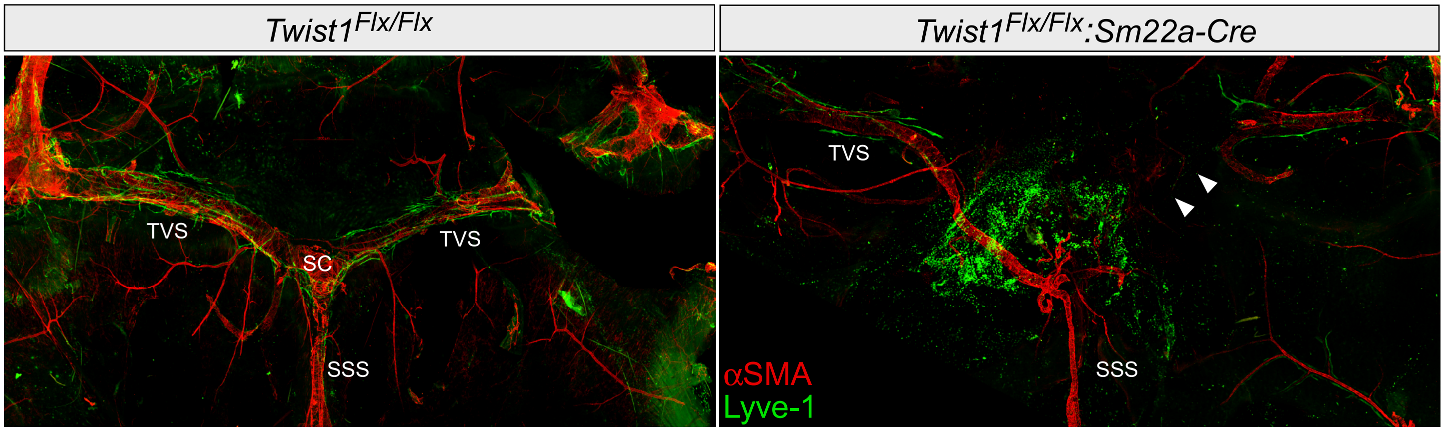

### Supplemental Figure 3

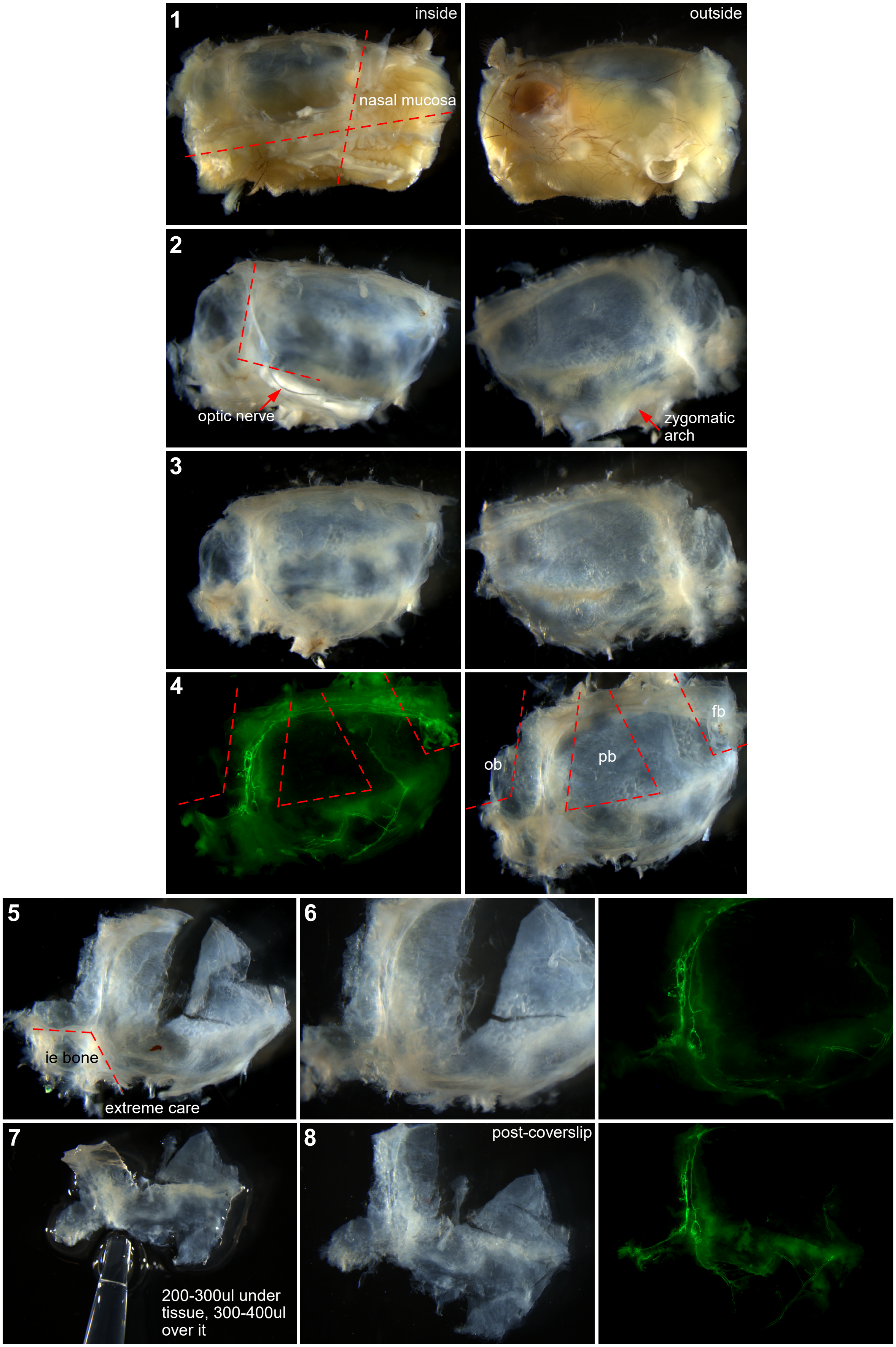

### Supplemental Figure 4

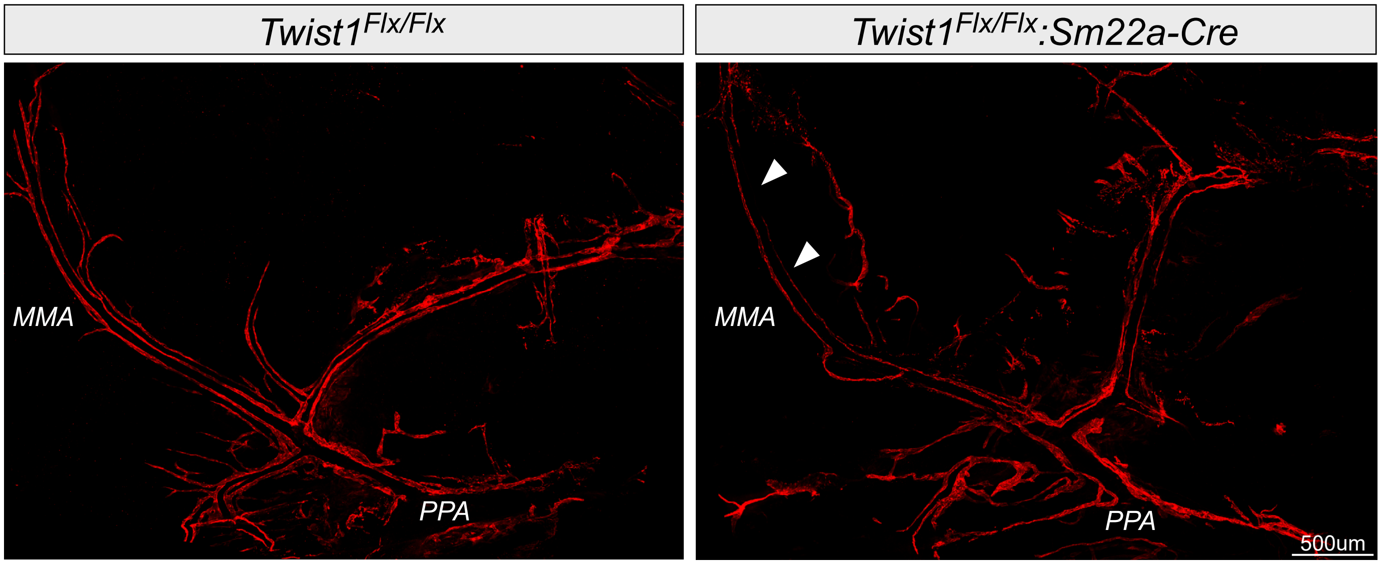
